# Decision-making in serial crystallography: a simple test to quickly determine whether sufficient data have been collected

**DOI:** 10.1101/2025.08.12.669835

**Authors:** David von Stetten, Arwen R Pearson

**Affiliations:** European Molecular Biology Laboratory (EMBL), Notkestraße 85, 22760 Hamburg, Germany; University of Hamburg, Institute for Nanostructure and Solid State Physics, Hamburg Centre for Ultrafast Imaging, HARBOR Bldg 610, Luruper Chaussee 149, 22761 Hamburg, Germany

**Keywords:** Serial crystallography, time-resolved crystallography, macromolecular crystallography, signal-to-noise

## Abstract

In standard rotational data collection for macromolecular crystallography data are normally collected from a single crystal, and the resulting data processing delivers metrics for data completeness and signal to noise that are well established. However, in serial crystallography it can be difficult to assess quickly whether enough data have been recorded to deliver a well scaled and complete dataset with sufficient signal to noise to address the scientific question being asked. Completeness alone is not an appropriate metric, as a nominally complete dataset can be obtained with a much smaller number of images, and thus multiplicity, than is needed to produce a final dataset with well estimated merged intensity values. Insufficient data result in alarmingly reasonable processing statistics and plausible electron density maps that contain almost no experimental signal, instead being dominated by the phases from the phasing model. We have therefore established a simple electron density-based test to determine whether enough data have been collected, and implemented this in the autoprocessing pipeline at the T-REXX endstation on beamline P14 at PETRA III. Importantly, the results of this test help guide decisions as to whether more data should be collected, or whether the experimenter can move onto a new time-point or sample.

**Synopsis:** We describe a simple test to determine whether sufficient data have been collected during a serial crystallographic experiment, and its incorporation into the autoprocessing pipeline at the T- REXX endstation on beamline P14 at the PETRA III synchrotron.

## 1. Introduction

Serial crystallography is a diffraction method in which a large number of single microcrystals are delivered to the probe beam in random orientations (Chapman *et al*., 2011). A number of sample delivery options are available, depending on source, beamline, and experiment type (Martiel *et al*., 2019). All serial experiments produce a series of diffraction patterns that are independent of each other in terms of orientation. During processing blank images are discarded and images containing diffraction spots, “hits”, are indexed and integrated. On the basis of the indexing, and associated orientation matrix that describes the alignment of each crystal with respect to the laboratory frame, the integrated reflections can be assembled or “ordered” into a 3D data set. As each diffraction pattern is from a different crystal and has been independently processed, the resulting dataset contains a spread of unit cell dimensions and varying intensity values for each reflection, for example as a result of varied crystal size, partiality, and/or varying incident beam intensity. Next, a scaling algorithm attempts to converge on a good estimate of the true intensity of each reflection before a final merging step. Scaling removes outliers, applies scaling factors on a per image basis, and, in more complex processing pathways, can attempt to model the partiality of reflections on each frame. Scaling of serial data fundamentally relies on the law of large numbers, making the assumption that if the population of intensity values for each reflection is sufficiently well sampled over a large number of independent measurements (*i*.*e*. individual diffraction patterns), the final mean value of the merged intensity will converge on the true value.

The danger in calculating electron density maps from datasets with insufficient sampling of each reflection is that the resulting maps are dominated by the model phases. This phase bias can result in seductively beautiful, but meaningless maps, leading the unwary crystallographer to assume the data are much better than they in fact are.

A key question during a serial diffraction experiment is therefore: when have sufficient data been collected to obtain a good enough estimate of the intensity of each reflection to allow confident interpretation of the resulting electron density maps? The relevant statistical metric here is the multiplicity. This describes, on average, how often each reflection has been observed. However, this aggregate metric can be misleading, for example in cases where there is a strong preferential orientation of the individual crystals and some reflections have been measured many times, but others much less frequently. Other metrics usually used to assess data quality are less useful, or in the case of completeness, actively misleading. A number of rules-of-thumb circulate in the serial community which attempt to help users decide when enough is enough in terms of data, but these can be effectively summarised as “more is always better”. This is not a particularly helpful guideline for decision making on the fly during data collection. We have therefore developed an electron density-based test that can be run as part of the autoprocessing pipeline during serial experiments. This provides rapid feedback to users as to whether they have reached sufficient multiplicity, and thus sufficient signal to noise, to have overcome model phase bias. For time-resolved experiments, it can help to indicate whether the expected structural changes are likely to be visible in the electron density map, or whether additional data are needed.

## 2. Methods

### 2.1 Summary of the current T-REXX autoprocessing pipeline

T-REXX (“time-resolved X-ray crystallography”) is an endstation on beamline P14 at the PETRA III synchrotron (DESY, Hamburg) dedicated to serial crystallographic experiments (von Stetten *et al*., 2019). The autoprocessing pipeline implemented on T-REXX takes the form of a bash script generated by MXCuBE (Gabadinho *et al*., 2010; Oscarsson *et al*., 2019) on the basis of user supplied information (starting model, cell & space group parameters) and the type of serial experiment selected (static structure, burst mode, HARE data collection, etc.). The script runs CrystFEL for peakfinding (*zaef*) (White *et al*., 2012), indexing (*XGandalf*) (Gevorkov *et al*., 2019), integration, scaling and merging (*partialator)* (White *et al*., 2016). *ambigator* is run automatically for those spacegroups where indexing ambiguities can occur (White *et al*., 2016). Once scaling and merging are complete, the resulting merged intensities are written to MTZ format and passed to the DIMPLE pipeline (Wojdyr *et al*., 2013), along with a user provided model (PDB format). If there is a mismatch in space group or large difference in cell parameters from the user provided model file, DIMPLE carries out molecular replacement using PHASER (McCoy *et al*., 2007). DIMPLE finishes with 4 cycles of jelly body and 8 cycles of restrained refinement using REFMAC5 (Murshudov *et al*., 2011) with TWIN refinement used in order to detect possible indexing ambiguities, that are not successfully resolved by *ambigator*. FFT from the CCP4 suite (Agirre *et al*., 2023) is then run to create 2*m*F_o_-*D*F_c_ (FWT and PHWT) and *m*F_o_-*D*F_c_ (DELFWT and PHDELWT) electron density maps and these are passed to the ISPyB database (Delagenière *et al*., 2011), together with the mtz and pdb file output by REFMAC5, additional data quality plots and a “Table 1” style summary of data processing statistics. Users can then view these files and the resulting electron density map and model through the EXI web interface (https://exi.embl-hamburg.de) using UGLYMOL (Wojdyr, 2017).

**Table 1.** Summary of test datasets used to demonstrate the PMRDD test.

| Protein | Endstation Source | Sample delivery Method | Resolution | Space group | # of crystals per test dataset* |
| --- | --- | --- | --- | --- | --- |
| Xylose isomerase<sup>1</sup> | T-REXX<br>PETRA III | HARE Chip<br>(fixed target) | 1.7 Å | I222 | 100, 310, 1000, 3000,<br>10001, 23141 |
| CTXM<sup>1</sup> (+<br>ambigator) | T-REXX<br>PETRA III | HARE Chip<br>(fixed target) | 1.7 Å | P3<sub>2</sub>21 | 102, 303, 1010, 3005,<br>10026, 27408 |
| CTXM<sup>1</sup> (-<br>ambigator) | T-REXX<br>PETRA III | HARE Chip<br>(fixed target) | 1.7 Å | P3<sub>2</sub>21 | 101, 297, 1008, 3000,<br>10009, 27421 |
| ADC<sup>2</sup> | MASSIF-3<br>ESRF | 3DMixD<br>(microfluidic chip) | 1.8 Å | P6<sub>1</sub>22 | 100, 300, 1000, 3000,<br>10000, 30000 |
| phyA<sup>3</sup> | SPB/SFX<br>EuXFEL | GDVN liquid jet | 2.2 Å | P2<sub>1</sub> | 101, 301, 1003, 3000,<br>10000, 30001 |
| OaPAC<sup>4</sup> | Cristallina<br>SwissFEL | MISP Chip<br>(fixed target) | 2.2 Å | P2<sub>1</sub>2<sub>1</sub>2 | 101, 302, 1001, 3001,<br>10000, 30000 |

\*The number of images per test dataset is not completely uniform due to the way in which the *partialator* programme within *CrystFEL* rejects images during the scale and merge step. <sup>1</sup>The xylose isomerase data and CTXM beta lactamase data have been previously reported in (Schulz *et al.*, 2025). <sup>2</sup>The aspartate decarboxylase data have been previously reported in (Monteiro *et al.*, 2020). <sup>3</sup>The phytochrome A data have been previously reported in (Nagano *et al.*, 2025). <sup>4</sup>The photoactivated adenylate cyclase data are unpublished and used here by kind permission of Sofia Kapetanaki.

### 2.2 Model preparation for the Perturbed Model Real-space Difference Density test (PMRDD)

To enable use of the PMRDD test in the T-REXX autoprocessing pipeline, prior to loading into MXCuBE, the user-supplied model is manually edited to remove a core, well ordered aromatic residue side chain that, if possible, is not close to the region of interest, *e*.*g*., the active site or ligand binding site. Additionally, a second core aromatic is changed to a different rotamer that ideally does not clash with the rest of the model.

The user supplied model is normally a refined model from a previous room temperature (serial or single crystal) or cryo experiment on the same macromolecular target, or a sufficiently similar model from the wwPDB. Due to the nature of the experiments on T-REXX (*i*.*e*. time-resolved serial measurements), T-REXX users, so far, have never lacked a good starting model, but in principle this test is usable for serial experiments where the goal is *de novo* structure determination via molecular replacement.

### 2.3. Generation of test datasets to demonstrate utility of the PMRDD test

To demonstrate the impact of increasing multiplicity on electron density map quality and to highlight the utility of the PMRDD test, and its robustness towards data collected in different kinds of serial experiments, we used several serial datasets collected using fixed targets at T-REXX (PETRA III) (von Stetten *et al*., 2019; Mehrabi *et al*., 2020) and Cristallina (SwissFEL) (Carrillo *et al*., 2023), a microfluidic device at MASSIF-3 (ESRF) (Von Stetten *et al*., 2020; Monteiro *et al*., 2020), and liquid jets at SPB/SFX (European XFEL) (Mancuso *et al*., 2019; Schulz *et al*., 2019) (Table 1). These were all processed using the standard T-REXX autoprocessing pipeline (identical processing parameters, aside from the XFEL datasets which were introduced into the T-REXX pipeline post integration) using CrystFEL v0.10.2 with increasing numbers of images included in each merged dataset. For the CTXM dataset, test datasets were processed with and without the use of *ambigator* to resolve indexing ambiguities. Full processing statistics are provided as supplementary data (Tables S1-S6).

## 3. Results

In all examples, the PMRDD test shows clearly when sufficient data have been included to overcome model phase bias (Figures 1-5 plus supplementary figures S1 and S2). This can be seen in the progressive appearance of positive difference electron density at the correct position for the moved and deleted residues, as well as negative difference density at the position of the moved residue.

**Figure 1.**
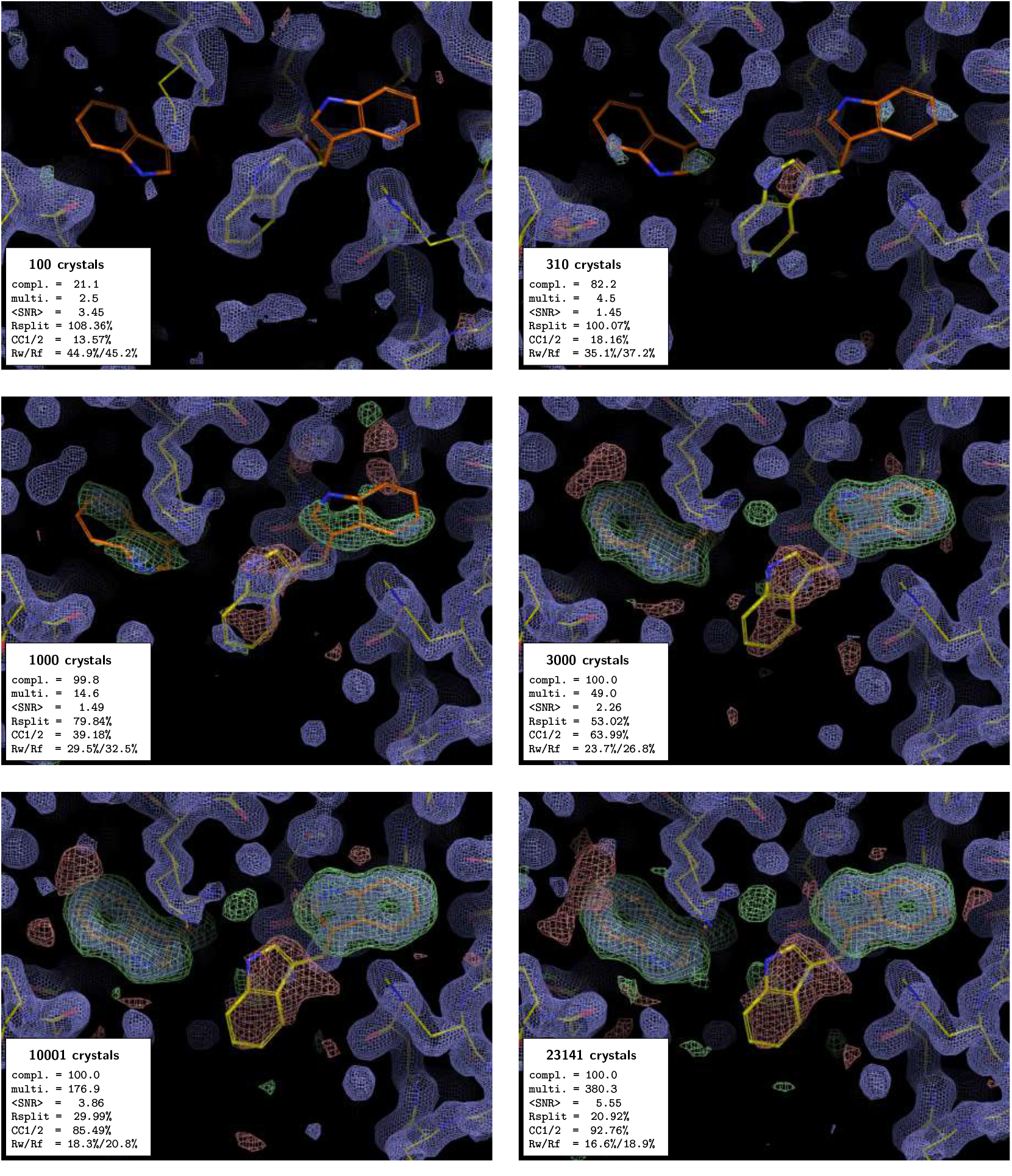
PMRDD test applied to a xylose isomerase serial dataset collected at T-REXX. Each panel shows a Pymol (v2.5.0) (Schrödinger, LLC, 2015) generated equivalent of a COOT (Emsley *et al*., 2010) screenshot, showing 2*m*F_o_-*D*F_c_ electron density maps contoured at 1.5 r.m.s.d. (blue mesh) and *m*F_o_-*D*F_c_ electron density maps contoured at ± 3.0 r.m.s.d. (green + 3.0 r.m.s.d. and red -3.0 r.m.s.d. mesh). In each panel, the “true” starting model is shown as sticks with carbons coloured orange. The edited “PMRDD model” is shown as sticks with carbons coloured yellow. In each panel, an inset box indicates the number of diffraction patterns included in the merge (number of crystals) and summarises the key data quality metrics (completeness [%], multiplicity, signal-to-noise ratio, R_split_ [%](White *et al*., 2012), CC1/2 [%] (Assmann *et al*., 2016), and overall R_work_ and R_free_ [%]). A complete table of data quality statistics can be found in the supplementary data Table S1.

**Figure 2.**
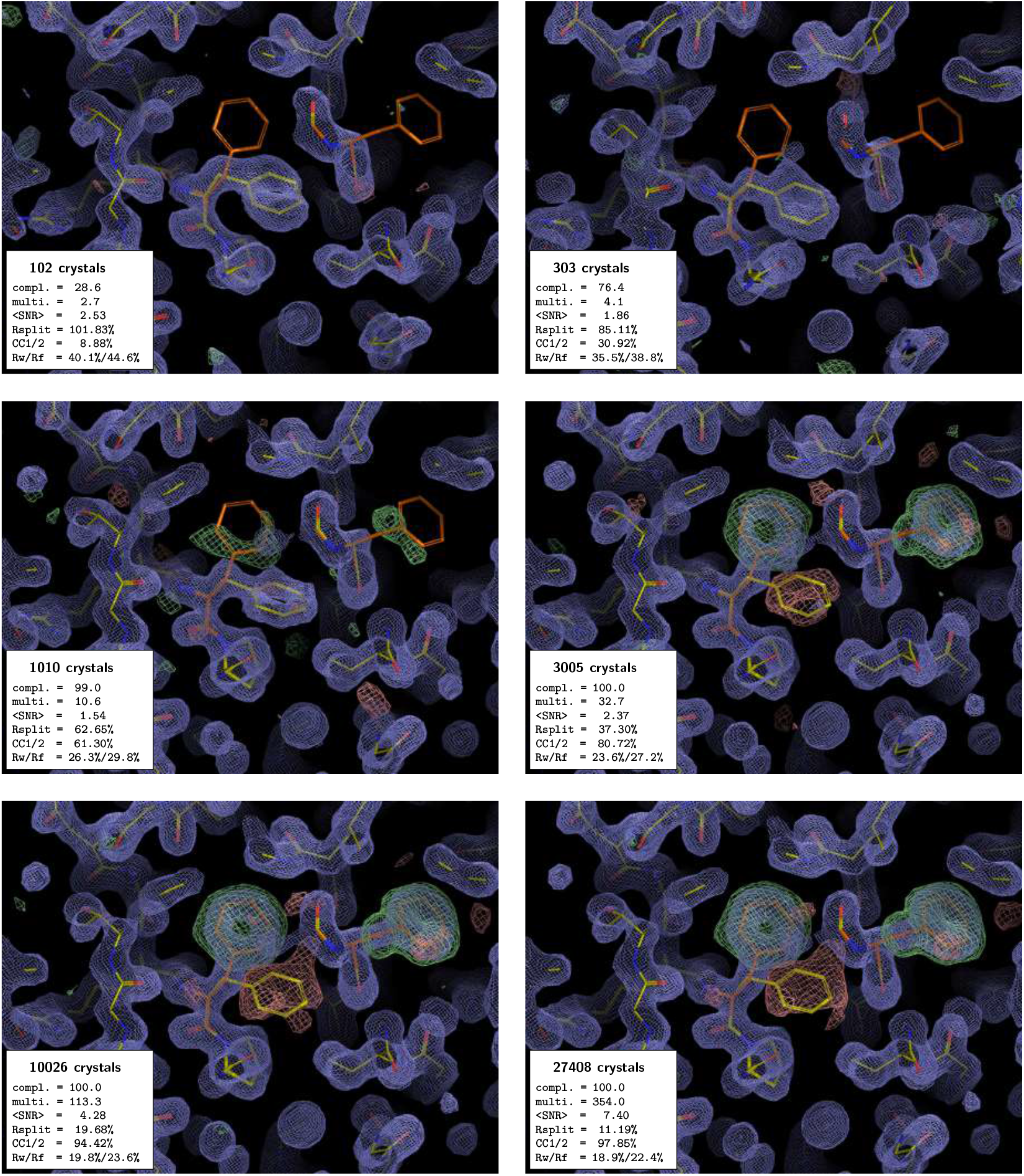
PMRDD test applied to a CTXM beta lactamase serial dataset collected at T-REXX, with the use of *ambigator* during the data processing. The panels, electron density maps and colouring are all as in Figure 1. A complete table of data quality statistics can be found in the supplementary data Table S2.

**Figure 3.**
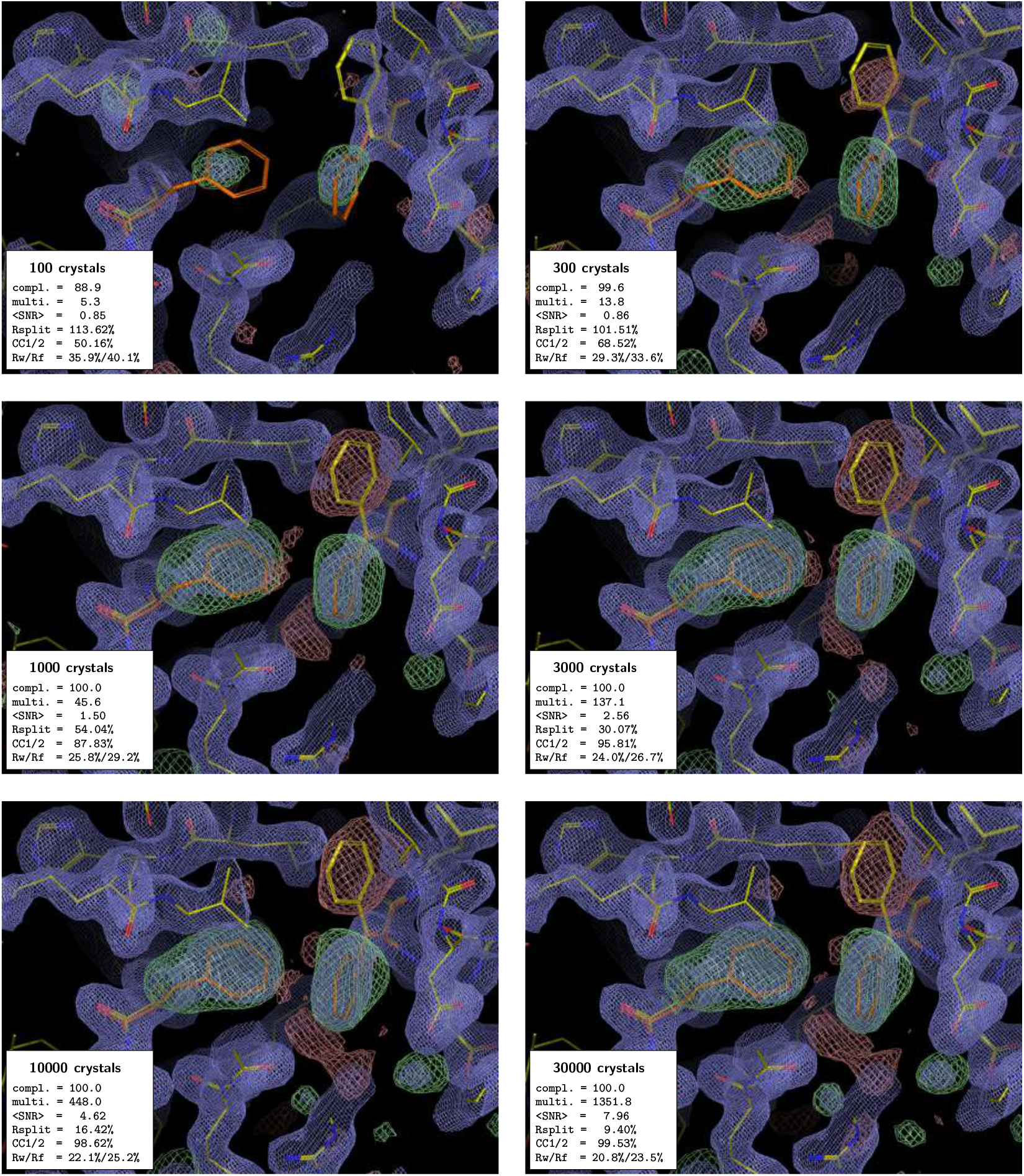
PMRDD test applied to an aspartate decarboxylase (ADC) serial dataset collected at MASSIF-3, ESRF. The panels, electron density maps and colouring are all as in Figure 1. A complete table of data quality statistics can be found in the supplementary data Table S3.

**Figure 4.**
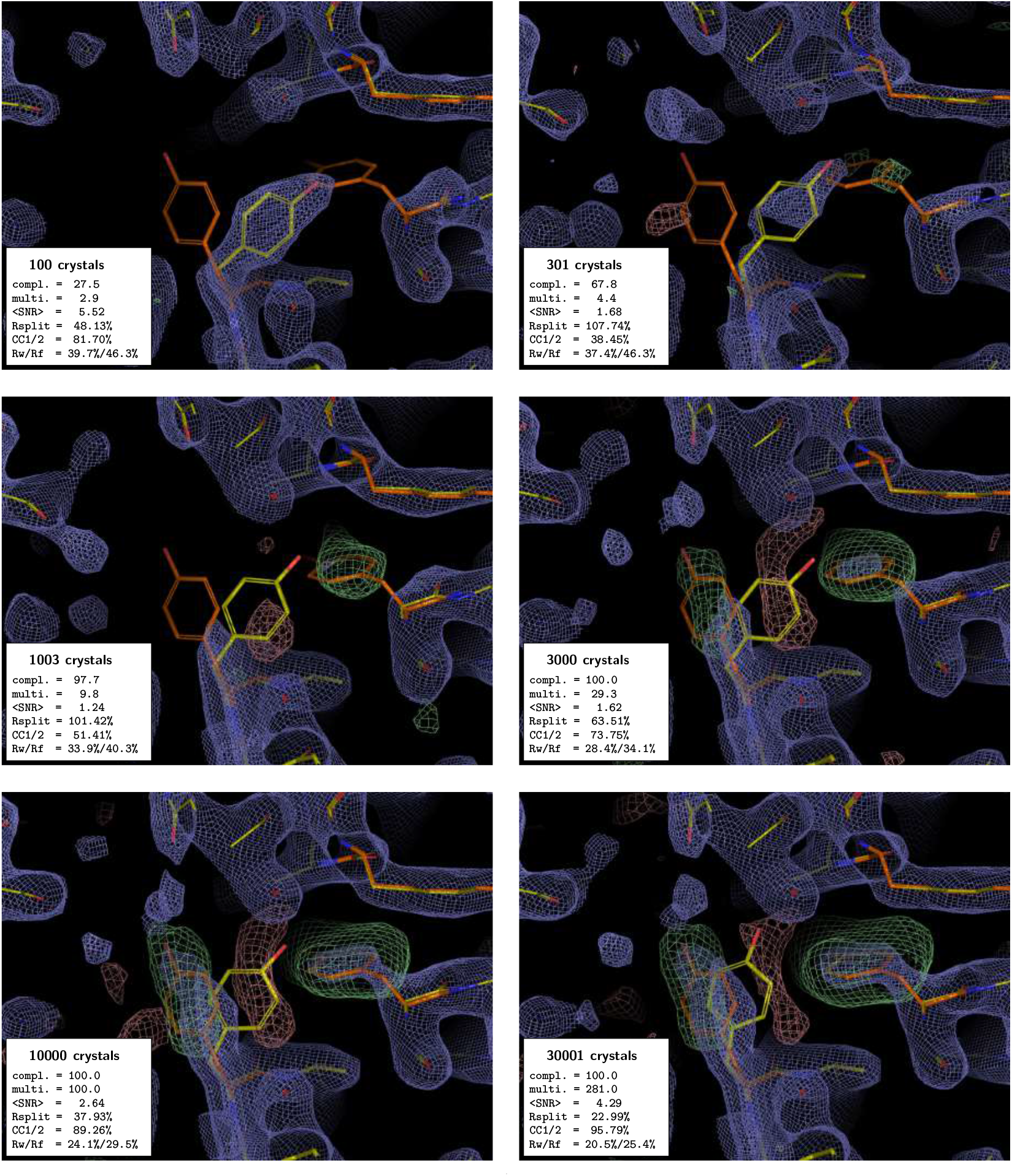
PMRDD test applied to a phytochrome A (phyA) serial dataset collected at the European XFEL. The panels, electron density maps and colouring are all as in Figure 1. A complete table of data quality statistics can be found in the supplementary data Table S4.

**Figure 5.**
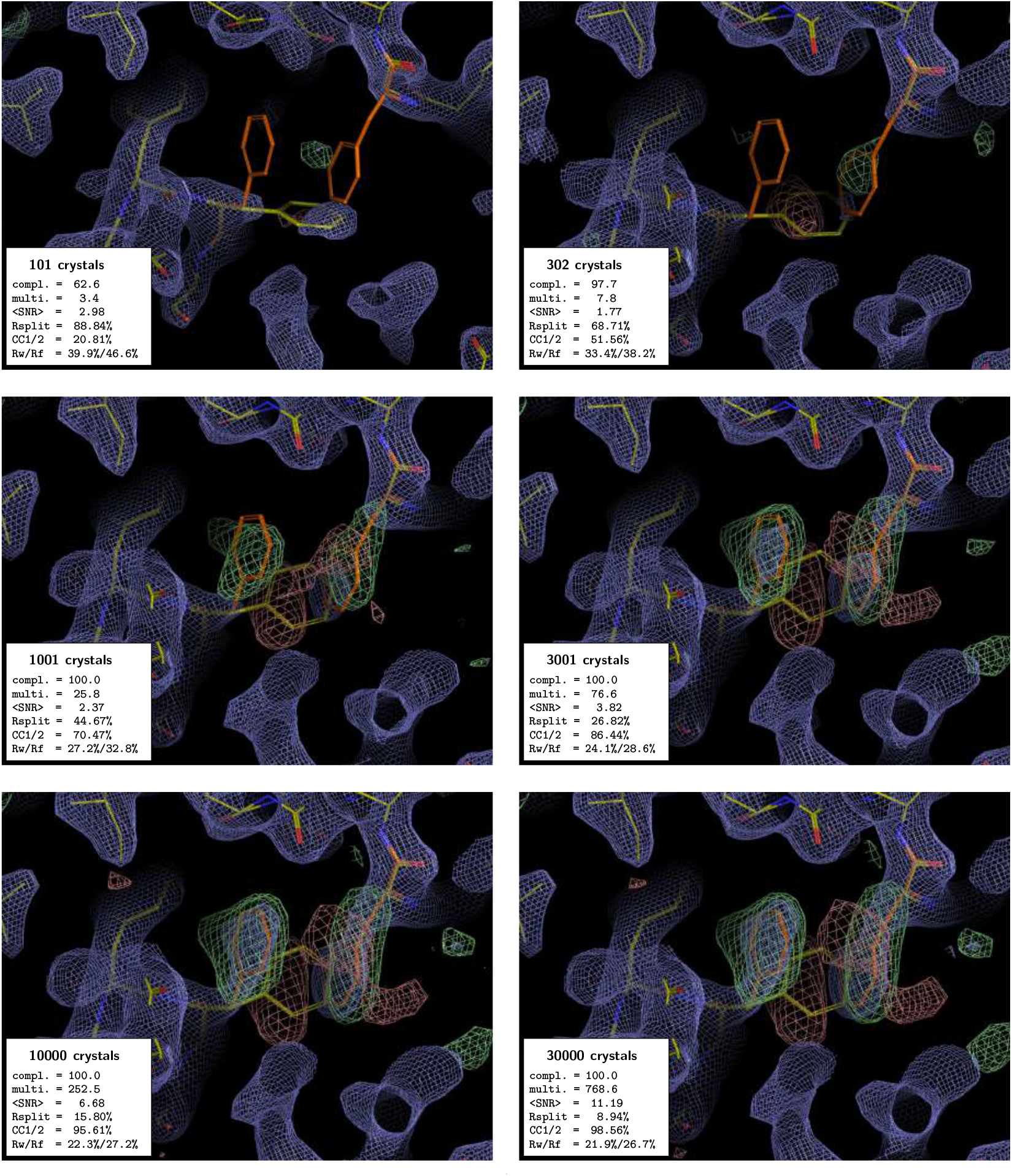
PMRDD test applied to a photoactivated adenylate kinase (OaPAC) dataset collected at Cristallina, SwissFEL. The panels, electron density maps and colouring are all as in Figure 1. A complete table of data quality statistics can be found in the supplementary data Table S5.

For the beginning crystallographer, we emphasise the fact that, even for datasets derived from a clearly insufficient number of diffraction patterns, the 2*m*F_o_-*D*F_c_ electron density maps appear to be surprisingly good. However, this is entirely due to model bias, where, due to the weakness of the experimental data, the 2*m*F_o_-*D*F_c_ map effectively becomes an F_c_ map. This is clearly evident in the maps calculated with low numbers of difraction patterns where 2*m*F_o_-*D*F_c_ electron density is present at the position of the residue that was moved to an incorrect position for the PMRDD test, and absent for the residue that was deleted. Only once the number of diffraction patterns in a dataset is sufficient, do the electron densities for these perturned residues show up correctly in both the 2*m*F_o_-*D*F_c_ and *m*F_o_-*D*F_c_ maps.

The test works equally well, regardless of resolution and space group. In all cases it can be seen that the data are technically complete long before there is clear difference density overcoming the model bias.

A few interesting additional observations can be made from these test data. First that the use of *ambigator* to resolve indexing ambiguities has a noticeable positive effect on the quality of the difference density, even though for the CTXM datasets with up to 1000 diffraction patterns *ambigator* fails to resolve the indexing ambiguity, and consequently *REFMAC5* determines these datasets as “twinned” (compare Figures 2 and S1 and Tables S2 and S6). However, for some datasets with a low number of crystals (e.g., 100- and 300-crystal datasets of xylose isomerase), REFMAC5 reports multiple twin domains (despite the absence of an indexing ambiguity) due to the rather poor data. Second, the data recorded using the 3DMixD microfluidic chip show clear difference density for the correct structure at a much lower number of crystals than the data recorded using fixed targets at synchrotrons or the GDVN nozzles at European XFEL. This could be due to the possibility for the crystals to rotate during the X-ray exposure (∼5 ms residence time in the beam) as they flow through the microfluidic device. This would result in measuring more fully-recorded reflections, potentially leading to a more stable scaling and convergence to a good estimate of their intensities already at lower multiplicity. However, it is also possible that the high symmetry of ADC is contributing. Further work will be needed to distinguish these possibilities.

## 4. Summary

The PMRDD test is a robust and simple test to help users decide when sufficient data have been collected in a serial experiment. It is easy to implement, either as a standalone test by an individual user (by simply deleting/moving a well-ordered residue that is not expected to change position during the experiment prior to phasing and map calculation) or within a data processing pipeline at a synchrotron or XFEL facility by providing a PDB file that has been modified as described here. It is useful at all resolutions tested and, maybe most importantly, is conceptually easy for novice users to grasp.

## Acknowledgements

We thank our users and collaborators who have agreed to use of their data in this paper. The xylose isomerase and CTXM data were used by kind permission of Eike Schulz (University Clinic Eppendorf, Hamburg) and Pedram Mehrabi (University of Hamburg). The phyA data were used by kind permission of Jon Hughes and Soschichiro Nagano (Max Planck Institute for Medical Research, Heidelberg & Free University of Berlin). The OaPAC data were used by kind permission of Sofia Kapetanaki (University of Pecs). Funding for development and the construction of the T-REXX endstation was received from the BMBF (Verbundforschung, 05K16GU1 and 05K19GU1, 05K22GU6). T-REXX beamtime was granted in the EMBL BAGs MX-660 and MX-1008. MASSIF-3 beamtime was granted through through ESRF proposal No. MX1919, integrated into the Serial Synchrotron Crystallography BAG (MX1887/MX1991). SPB/SFX beamtime at European XFEL was awarded under proposal 4511. Cristallina beamtime at SwissFEL was awarded under proposal 20231124.

## Conflicts of interest

The authors declare no conflict of interest in this paper.

## Data availability

The xylose isomerase data and CTXM beta lactamase data have been previously reported in (Schulz *et al*., 2025, PDBs 9G5N & 9G7W respectively). The aspartate decarboxylase data have been previously reported in (Monteiro *et al*., 2020, PDB 6RXH), and the raw data are available at https://www.cxidb.org/id-147.html, dataset “chip3_ADC5_12_3mm.tgz”. The phytochrome A data have been previously reported in (Nagano *et al*., 2025, 9ER4). The OaPAC data are unpublished and used here by kind permission of Sofia Kapetanaki.

## Notes

### Competing Interest Statement

The authors have declared no competing interest.

