## Supplemental data tables and figures for "Decision-making in serial crystallography: a simple test to quickly determine whether sufficient data have been collected"

### Supporting information

**Table S1** Data processing statistics for the xylose isomerase dataset (Schulz *et al.*, 2025). Data were collected at the T-REXX endstation on EMBL beamline P14 at PETRA III. Data were collected at 295 K. Samples were delivered to the X-ray beam using HARE-Chips (Mehrabi *et al.*, 2020).

| **Data collection** | | | | | | |
|---|---|---|---|---|---|---|
| # diffraction patterns | 100 | 310 | 100 | 3000 | 10001 | 23141 |
| Space group | I222 | | | | | |
| Unit cell dimensions (Å, °) | 94.2, 103.0, 99.2; 90.0, 90.0, 90.0 | | | | | |
| Resolution range (Å) | 71.4 – 1.70 | 71.4 – 1.70 | 71.4 – 1.70 | 71.4 – 1.70 | 71.4 – 1.70 | 71.4 – 1.70 |
| | (1.76 – 1.70) | (1.76 – 1.70) | (1.76 – 1.70) | (1.76 – 1.70) | (1.76 – 1.70) | (1.76 – 1.70) |
| Wilson B factor (Å$^2$) | 20.0 | 21.4 | 20.4 | 20.2 | 20.6 | 20.2 |
| # measured reflections | 28305 | 197921 | 777629 | 2616262 | 9440232 | 20298653 |
| # unique reflections | 11285 | 43880 | 53241 | 53373 | 53373 | 53373 |
| Completeness (%) | 21.1 (9.7) | 82.2 (65.0) | 99.8 (99.2) | 100.0 (100.0) | 100.0 (100.0) | 100.0 (100.0) |
| Multiplicity | 2.5 (2.2) | 4.5 (3.1) | 14.6 (8.6) | 49.0 (28.9) | 176.9 (104.2) | 380.3 (224.3) |
| Signal-noise-ratio | 3.5 (1.1) | 1.4 (0.6) | 1.5 (0.6) | 2.3 (1.2) | 3.9 (2.2) | 5.5 (3.3) |
| Rsplit (%) | 108.4 (164.8) | 100.1 (207.8) | 79.8 (176.4) | 53.0 (86.8) | 30.0 (48.6) | 20.9 (34.0) |
| CC* (%) | 48.9 (NaN) | 55.4 (34.0) | 75.0 (45.9) | 88.3 (70.8) | 96.0 (88.2) | 98.1 (93.0) |
| CC1/2 (%) | 13.6 (45.8) | 18.2 (6.1) | 39.2 (11.8) | 64.0 (33.4) | 85.5 (63.6) | 92.8 (76.2) |

| | | | | | | |
|---|---|---|---|---|---|---|
| **Refinement** | | | | | | |
| Resolution range (Å) | 71.5 – 1.56 | 71.5 – 1.56 | 71.5 – 1.70 | 71.5 – 1.70 | 71.5 – 1.70 | 71.5 – 1.70 |
| Mean B (Å$^2$) | 10.7 | 12.5 | 14.5 | 15.1 | 15.5 | 15.2 |
| $R_{work}$ (%) | 44.9 | 35.1 | 29.5 | 23.7 | 18.3 | 16.6 |
| $R_{free}$ (%) | 45.2 | 37.2 | 32.5 | 26.8 | 20.8 | 18.9 |
| # Twin domains[2] | 6 | 4 | 0 | 0 | 0 | 0 |

[1] Numbers in parentheses refer to the highest resolution shell

[2] As discussed in §3 of the main text, twin domains are erroneously detected by REFMAC due to the poor data quality.

**Table S2** Data processing statistics for the CTXM beta-lactamase dataset using ambigator (Schulz *et al.*, 2025). Data were collected at the T-REXX endstation on EMBL beamline P14 at PETRA III. Data were collected at 295 K. Samples were delivered to the X-ray beam using HARE-Chips (Mehrabi *et al.*, 2020).

| | | | | | | |
|---|---|---|---|---|---|---|
| **Data collection** | | | | | | |
| # diffraction patterns | 102 | 303 | 1010 | 3005 | 10026 | 27408 |
| Space group | $P3_221$ | | | | | |
| Unit cell dimensions (Å, °) | 41.9, 41.9, 232.8; 90.0, 90.0, 120.0 | | | | | |
| Resolution range (Å) | 116.3 – 1.70 (1.76 – 1.70) | 116.3 – 1.70 (1.76 – 1.70) | 116.3 – 1.70 (1.76 – 1.70) | 116.3 – 1.70 (1.76 – 1.70) | 116.3 – 1.70 (1.76 – 1.70) | 116.3 – 1.70 (1.76 – 1.70) |
| Wilson B factor (Å$^2$) | 24.4 | 24.6 | 24.6 | 24.6 | 24.3 | 24.1 |
| # measured reflections | 20776 | 85778 | 289412 | 897047 | 3111476 | 9718004 |
| # unique reflections | 7838 | 20979 | 27178 | 27452 | 27454 | 27455 |
| Completeness (%) | 28.6 (15.2) | 76.4 (58.7) | 99.0 (97.5) | 100.0 (100.0) | 100.0 (100.0) | 100.0 (100.0) |
| Multiplicity | 2.7 (2.2) | 4.1 (3.0) | 10.6 (6.5) | 32.7 (19.7) | 113.3 (68.7) | 354.0 (214.2) |
| Signal-noise-ratio | 2.5 (0.1) | 1.9 (0.7) | 1.5 (0.4) | 2.4 (0.6) | 4.3 (1.1) | 7.4 (2.2) |
| Rsplit (%) | 101.8 (-266.0) | 85.1 (308.8) | 62.6 (313.5) | 37.3 (174.4) | 19.7 (90.7) | 11.2 (45.2) |
| CC* (%) | 40.4 (54.0) | 68.7 (NaN) | 87.2 (32.4) | 94.5 (50.3) | 98.6 (81.1) | 99.5 (94.8) |
| CC1/2 (%) | 8.9 (17.0) | 30.9 (0.6) | 61.3 (5.5) | 80.7 (14.5) | 94.4 (48.9) | 97.8 (81.5) |
| **Refinement** | | | | | | |

| | | | | | | |
|---|---|---|---|---|---|---|
| Resolution range (Å) | 77.6 – 1.70 | 77.6 – 1.70 | 77.6 – 1.70 | 77.6 – 1.70 | 77.6 – 1.70 | 77.6 – 1.70 |
| Mean B (Å$^2$) | 18.5 | 20.8 | 21.2 | 22.8 | 22.9 | 22.8 |
| $R_{work}$ (%) | 40.1 | 35.5 | 26.3 | 23.6 | 19.8 | 18.9 |
| $R_{free}$ (%) | 44.6 | 38.8 | 29.8 | 27.2 | 23.6 | 22.4 |
| # Twin domains[2] | 2 | 2 | 2 | 0 | 0 | 0 |

[1] Numbers in parentheses refer to the highest resolution shell

[2] As discussed in §3 of the main text, twin domains are erroneously detected by REFMAC when *ambigator* is unable to resolve the indexing ambiguity that arises during processing of a serial dataset.

**Table S3** Data processing statistics for the aspartate decarboxylase (ADC) dataset (Monteiro *et al.*, 2020). Data were collected at the MASSIF-3 beamline at ESRF. Data were collected at 295 K. Samples were delivered to the X-ray beam using the 3DMiXD microfluidic chip (Monteiro *et al.*, 2020).

| | | | | | | |
|---|---|---|---|---|---|---|
| **Data collection** | | | | | | |
| # diffraction patterns | 100 | 300 | 1000 | 3000 | 10000 | 30000 |
| Space group | $P6_122$ | | | | | |
| Unit cell dimensions (Å, °) | 72.0, 72.0, 216.8; 90.0, 90.0, 120.0 | | | | | |
| Resolution range (Å) | 108.7 – 1.80 (1.86 – 1.80) | 108.7 – 1.80 (1.86 – 1.80) | 108.7 – 1.80 (1.86 – 1.80) | 108.7 – 1.80 (1.86 – 1.80) | 108.7 – 1.80 (1.86 – 1.80) | 108.7 – 1.80 (1.86 – 1.80) |
| Wilson B factor ($Å^2$) | 26.8 | 27.5 | 27.4 | 27.2 | 27.3 | 26.9 |
| # measured reflections | 150022 | 440087 | 1457661 | 4384787 | 14327015 | 43227855 |
| # unique reflections | 28438 | 31860 | 31971 | 31979 | 31979 | 31979 |
| Completeness (%) | 88.9 (76.3) | 99.6 (99.1) | 100.0 (100.0) | 100.0 (100.0) | 100.0 (100.0) | 100.0 (100.0) |
| Multiplicity | 5.3 (3.6) | 13.8 (8.5) | 45.6 (27.7) | 137.1 (83.8) | 448.0 (274.1) | 1351.8 (827.7) |
| Signal-noise-ratio | 0.8 (0.1) | 0.9 (0.1) | 1.5 (0.1) | 2.6 (0.3) | 4.6 (0.6) | 8.0 (1.0) |
| Rsplit (%) | 113.6 (14991.0) | 101.5 (1285.3) | 54.0 (739.1) | 30.1 (367.7) | 16.4 (204.2) | 9.4 (111.1) |
| CC* (%) | 81.7 (44.1) | 90.2 (17.7) | 96.7 (20.6) | 98.9 (33.2) | 99.7 (52.7) | 99.9 (76.4) |
| CC1/2 (%) | 50.2 (10.8) | 68.5 (1.6) | 87.8 (2.2) | 95.8 (5.8) | 98.6 (16.1) | 99.5 (41.1) |
| **Refinement** | | | | | | |
| Resolution range (Å) | 62.4 – 1.80 | 62.4 – 1.80 | 62.4 – 1.80 | 62.4 – 1.80 | 62.4 – 1.80 | 54.1 – 1.80 |

| | | | | | | |
|---|---|---|---|---|---|---|
| Mean B ($Å^2$) | 21.4 | 30.6 | 33.5 | 31.4 | 30.3 | 29.6 |
| $R_{work}$ (%) | 35.9 | 29.3 | 25.8 | 24.0 | 22.1 | 20.8 |
| $R_{free}$ (%) | 40.1 | 33.6 | 29.2 | 26.7 | 25.2 | 23.5 |

[1] Numbers in parentheses refer to the highest resolution shell

**Table S4** Data processing statistics for the phytochrome A (phyA) dataset (Nagano *et al.*, 2025). Data were collected at the SPB/SFX instrument at European XFEL. Data were collected at 295 K. Samples were delivered to the X-ray beam using a GDVN (Schulz *et al.*, 2019).

| | | | | | | |
|---|---|---|---|---|---|---|
| **Data collection** | | | | | | |
| # diffraction patterns | 100 | 301 | 1003 | 3000 | 10000 | 30001 |
| Space group | | | $P2_1$ | | | |
| Unit cell dimensions (Å, °) | | | 56.5, 115.0, 69.8; 90.0, 92.7, 90.0 | | | |
| Resolution range (Å) | 16.9 – 2.20 (2.28 – 2.20)[1] | 18.5 – 2.20 (2.28 – 2.20) | 20.3 – 2.20 (2.28 – 2.20) | 20.5 – 2.20 (2.28 – 2.20) | 20.5 – 2.20 (2.28 – 2.20) | 20.5 – 2.20 (2.28 – 2.20) |
| Wilson B factor ($Å^2$) | -18.2 | 16.1 | 44.8 | 39.8 | 40.7 | 41.3 |
| # measured reflections | 36315 | 133496 | 431158 | 1322720 | 4512693 | 12680322 |
| # unique reflections | 12421 | 30590 | 44102 | 45124 | 45132 | 45132 |
| Completeness (%) | 27.5 (21.7) | 67.8 (59.0) | 97.7 (96.5) | 100 (100) | 100 (100) | 100 (100) |
| Multiplicity | 2.9 (2.7) | 4.4 (3.7) | 9.8 (7.5) | 29.3 (22.1) | 100.0 (75.1) | 281.0 (212.3) |
| Signal-noise-ratio | 5.5 (1.6) | 1.7 (0.0) | 1.2 (0.4) | 1.6 (0.3) | 2.6 (0.5) | 4.3 (0.8) |
| Rsplit (%) | 48.1 (165.8) | 107.7 (-437.0) | 101.4 (772.9) | 63.5 (414.9) | 37.9 (200.2) | 23.0 (125.2) |
| CC* (%) | 94.8 (39.8) | 74.5 (39.0) | 82.4 (52.7) | 92.1 (55.5) | 97.1 (73.8) | 98.9 (78.1) |
| CC1/2 (%) | 81.7 (8.6) | 38.5 (8.2) | 51.4 (16.1) | 73.8 (18.2) | 89.3 (37.5) | 95.8 (43.8) |
| **Refinement** | | | | | | |
| Resolution range (Å) | 20.5 – 2.11 | 19.9 – 2.14 | 20.3 – 2.20 | 20.5 – 2.20 | 20.5 – 2.20 | 20.5 – 2.20 |
| Mean B ($Å^2$) | 30.2 | 36.1 | 44.8 | 45.3 | 46.7 | 45.8 |

| | | | | | | |
|---|---|---|---|---|---|---|
| $R_{work}$ (%) | 39.7 | 37.4 | 33.9 | 28.4 | 24.1 | 20.5 |
| $R_{free}$ (%) | 46.3 | 46.3 | 40.3 | 34.1 | 29.5 | 25.4 |
| # Twin domains[2] | 2 | 2 | 0 | 0 | 0 | 0 |

[1] Numbers in parentheses refer to the highest resolution shell

[2] As discussed in §3 of the main text, twin domains are erroneously detected by REFMAC due to the poor data quality.

**Table S5** Data processing statistics for the photoactivated adenylate cyclase (OaPAC) dataset. Data were collected at the Cristallina endstation on the ARAMIS-ALVRA endstation at SwissFEL. Data were collected at 295 K. Samples were delivered to the X-ray beam using MISP-Chips (Carrillo *et al.*, 2023).

| | | | | | | |
|---|---|---|---|---|---|---|
| **Data collection** | | | | | | |
| # diffraction patterns | 101 | 302 | 1001 | 3001 | 10000 | 30000 |
| Space group | $P2_12_12$ | | | | | |
| Unit cell dimensions (Å, °) | 103.0, 55.0, 73.0; 90.0, 90.0, 90.0 | | | | | |
| Resolution range (Å) | 42.0 – 2.20 (2.28 – 2.20) | 43.9 – 2.20 (2.28 – 2.20) | 43.9 – 2.20 (2.28 – 2.20) | 43.9 – 2.20 (2.28 – 2.20) | 43.9 – 2.20 (2.28 – 2.20) | 43.9 – 2.20 (2.28 – 2.20) |
| Wilson B factor (Å$^2$) | 40.3 | 37.2 | 35.4 | 34.8 | 34.7 | 35.3 |
| # measured reflections | 46161 | 165193 | 560286 | 1665912 | 5491412 | 16719231 |
| # unique reflections | 13616 | 21254 | 21750 | 21752 | 21752 | 21752 |
| Completeness (%) | 62.6 (48.4) | 97.7 (95.8) | 100.0 (100.0) | 100.0 (100.0) | 100.0 (100.0) | 100.0 (100.0) |
| Multiplicity | 3.4 (2.7) | 7.8 (5.5) | 25.8 (18.0) | 76.6 (53.5) | 252.5 (176.6) | 768.6 (537.3) |
| Signal-noise-ratio | 3.0 (1.2) | 1.8 (0.8) | 2.4 (1.1) | 3.8 (1.9) | 6.7 (3.4) | 11.2 (5.8) |
| Rsplit (%) | 88.8 (-6579.5) | 68.7 (160.5) | 44.7 (95.8) | 26.8 (52.2) | 15.8 (28.1) | 8.9 (16.4) |
| CC* (%) | 58.7 (48.1) | 82.5 (59.4) | 90.9 (74.5) | 96.3 (90.7) | 98.9 (96.9) | 99.6 (98.9) |
| CC1/2 (%) | 20.8 (13.1) | 51.6 (21.4) | 70.5 (38.5) | 86.4 (69.8) | 95.6 (88.4) | 98.6 (95.7) |
| **Refinement** | | | | | | |

| | | | | | | |
|---|---|---|---|---|---|---|
| Resolution range (Å) | 43.9 – 2.20 | 43.9 – 2.20 | 43.9 – 2.20 | 43.9 – 2.20 | 43.9 – 2.20 | 43.9 – 2.20 |
| Mean B ($Å^2$) | 38.7 | 43.0 | 45.4 | 47.0 | 47.4 | 47.7 |
| $R_{work}$ (%) | 39.9 | 33.4 | 27.2 | 24.1 | 22.3 | 21.9 |
| $R_{free}$ (%) | 46.6 | 38.2 | 32.8 | 28.6 | 27.2 | 26.7 |

[1] Numbers in parentheses refer to the highest resolution shell

**Table S6** Data processing statistics for the CTXM beta-lactamase dataset, processed without ambigator. Data were collected at the T-REXX endstation on EMBL beamline P14 at PETRA III. Data were collected at 295 K. Samples were delivered to the X-ray beam using HARE-Chips (Mehrabi *et al.*, 2020).

| | | | | | | |
|---|---|---|---|---|---|---|
| **Data collection** | | | | | | |
| # diffraction patterns | 101 | 297 | 1008 | 3000 | 10009 | 27421 |
| Space group | P3221 | | | | | |
| Unit cell dimensions (Å, °) | 41.9, 41.9, 232.8; 90.0, 90.0, 120.0 | | | | | |
| Resolution range (Å) | 116.3 – 1.70 (1.76 – 1.70) | 116.3 – 1.70 (1.76 – 1.70) | 116.3 – 1.70 (1.76 – 1.70) | 116.3 – 1.70 (1.76 – 1.70) | 116.3 – 1.70 (1.76 – 1.70) | 116.3 – 1.70 (1.76 – 1.70) |
| Wilson B factor ($Å^2$) | 18.6 | 25.8 | 24.5 | 24.7 | 24.4 | 24.2 |
| # measured reflections | 20412 | 84145 | 288693 | 893643 | 3106336 | 9720847 |
| # unique reflections | 7778 | 20809 | 27178 | 27451 | 27454 | 27455 |
| Completeness (%) | 28.3 (14.5) | 75.8 (57.8) | 99.0 (97.5) | 100.0 (100.0) | 100.0 (100.0) | 100.0 (100.0) |
| Multiplicity | 2.6 (2.3) | 4.0 (2.9) | 10.6 (6.5) | 32.6 (19.7) | 113.1 (68.6) | 354.1 (214.3) |
| Signal-noise-ratio | 2.4 (0.2) | 1.6 (0.5) | 1.5 (0.7) | 2.4 (0.6) | 4.2 (1.1) | 7.3 (2.2) |
| Rsplit (%) | 98.9 (89.6) | 84.4 (314.3) | 63.5 (307.1) | 39.8 (182.6) | 22.1 (94.0) | 12.8 (46.6) |
| CC* (%) | 49.8 (NaN) | 68.1 (38.1) | 86.2 (33.6) | 93.6 (46.1) | 97.8 (70.4) | 99.2 (91.0) |
| CC1/2 (%) | 14.2 (98.3) | 30.2 (7.8) | 59.2 (6.0) | 77.9 (11.9) | 91.8 (32.9) | 97.0 (70.7) |
| **Refinement** | | | | | | |
| Resolution range (Å) | 77.6 – 1.70 | 77.6 – 1.70 | 77.6 – 1.70 | 77.6 – 1.70 | 77.6 – 1.70 | 77.6 – 1.70 |

| | | | | | | |
|---|---|---|---|---|---|---|
| Mean B (Å$^2$) | 16.7 | 21.2 | 21.1 | 22.5 | 22.5 | 22.4 |
| $R_{work}$ (%) | 40.9 | 34.7 | 26.0 | 19.9 | 15.4 | 13.7 |
| $R_{free}$ (%) | 44.8 | 36.4 | 30.1 | 22.9 | 18.9 | 17.3 |
| # Twin domains[2] | 2 | 2 | 2 | 2 | 2 | 2 |

[1] Numbers in parentheses refer to the highest resolution shell

[2] As discussed in §3 of the main text, twin domains are erroneously detected by REFMAC throughout when *ambigator* is not used to resolve the indexing ambiguity.

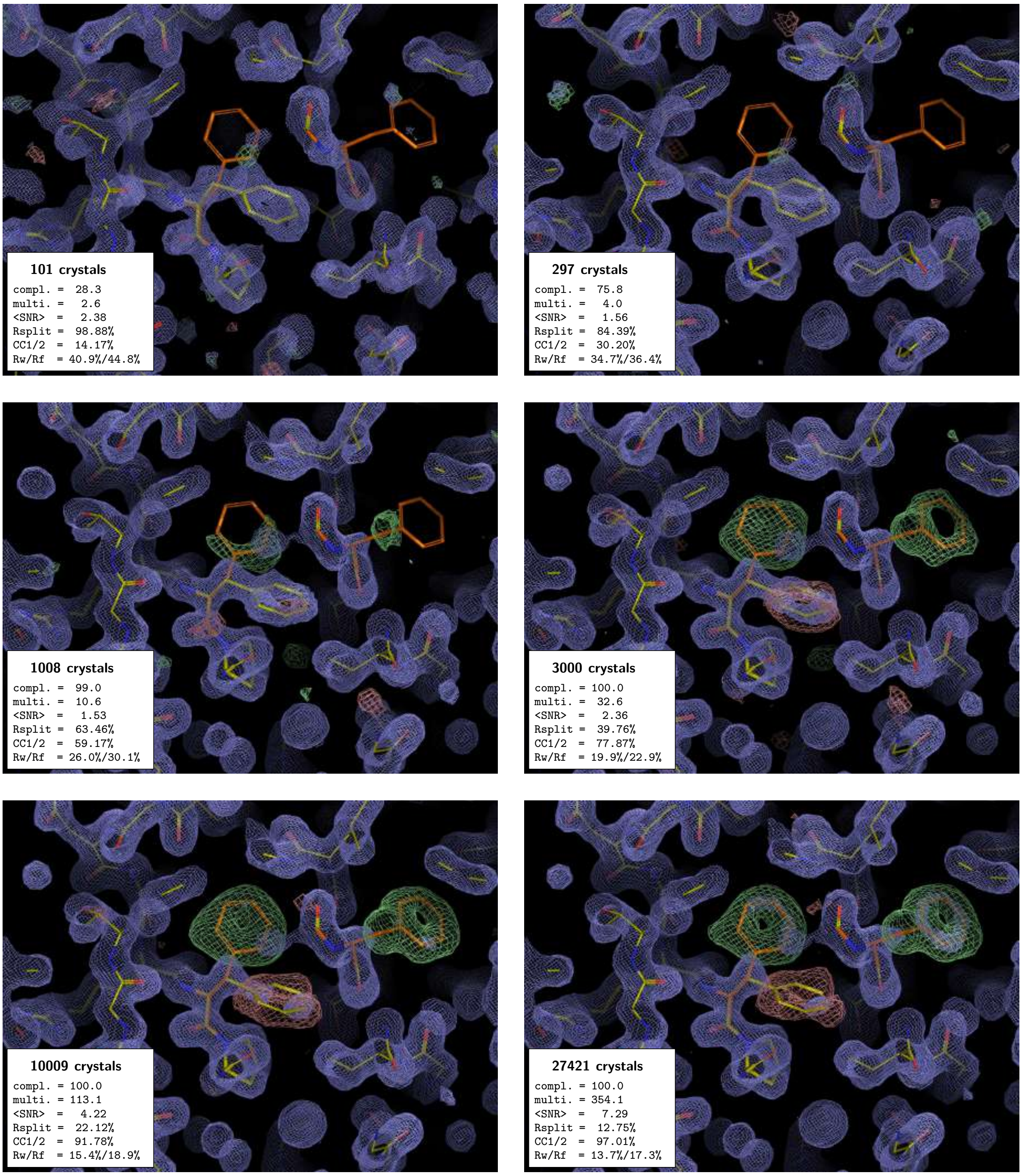


**Figure S1** PMRDD test applied to a CTXM beta lactamase serial dataset collected at T-REXX, without the use of *ambigator* during data processing. Each panel shows a Pymol (v2.5.0) (Schrödinger, LLC, 2015) generated equivalent of a COOT (Emsley *et al.*, 2010) screenshot, showing $2mF_o$-$DF_c$ electron density maps contoured at 1.5 r.m.s.d. (blue mesh) and $mF_o$-$DF_c$ electron density maps contoured at ± 3.0 r.m.s.d. (green + 3.0 r.m.s.d. and red -3.0 r.m.s.d. mesh). In each panel, the "true" starting model is shown as sticks with carbons coloured orange. The edited "PMRDD model" is shown as sticks with carbons coloured yellow. In each panel, an inset box indicates the number of diffraction patterns included in the merge (number of crystals) and summarises the key data quality metrics (completeness [%], multiplicity, signal-to-noise ratio, $R_{split}$ [%](White *et al.*, 2012), CC1/2

[%] (Assmann *et al.*, 2016), and overall $R_{work}$ and $R_{free}$ [%]). A complete table of data quality statistics can be found in the supplementary data Table S6.

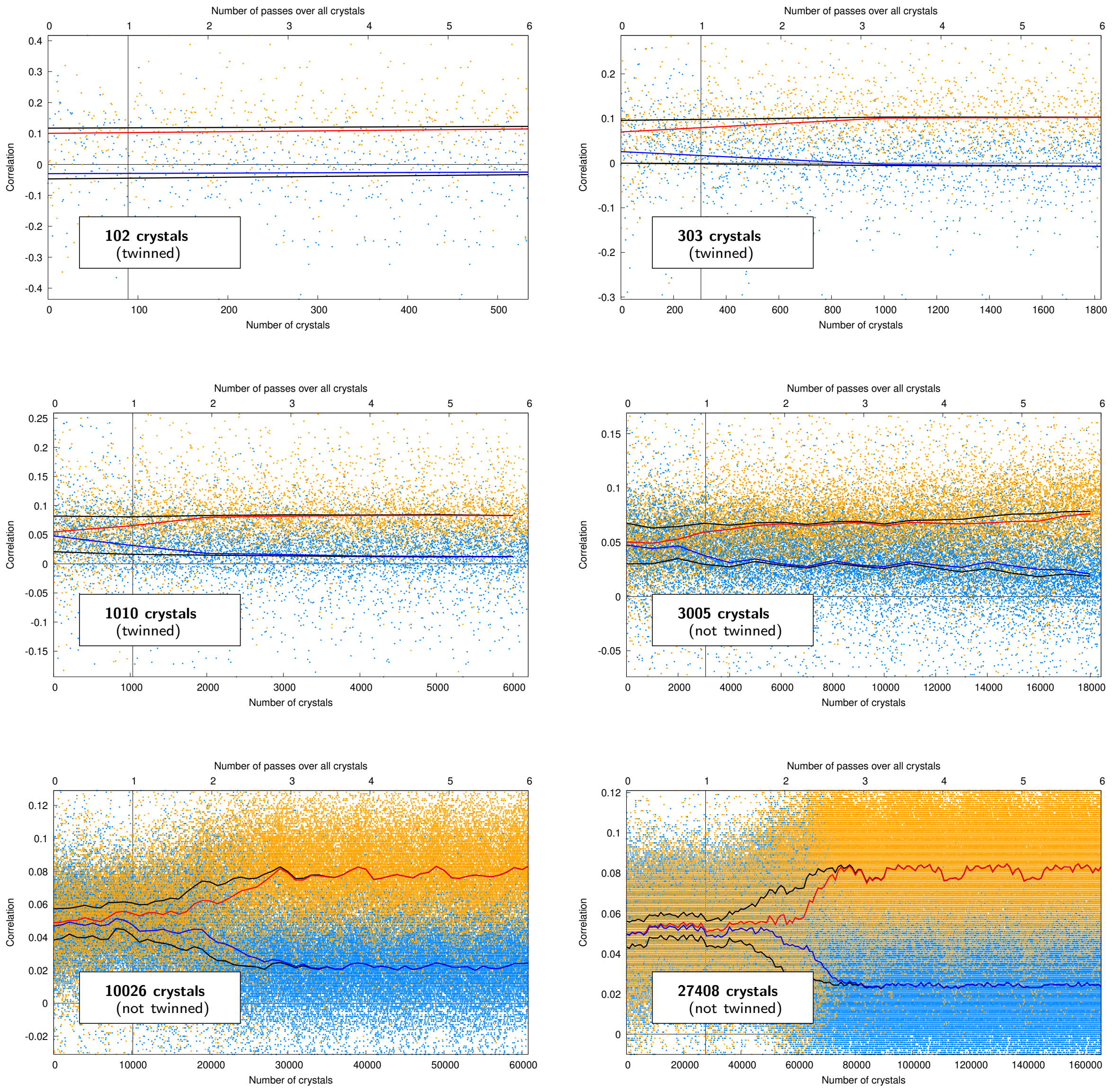


**Figure S2** *Ambigator* correlation plots for the CTXM beta lactamase serial dataset collected at T-REXX, corresponding to the electron density images shown in Figure 2. Each panel contains an inset indicating the number of diffraction patterns included in the merge (number of crystals) and indicates whether REFMAC identifies the data as twinned or not. At low numbers of crystals included in the merged dataset, the indexing ambiguity cannot be resolved by *ambigator* and REFMAC identifies the data as twinned. By ~3000 crystals in the merge *ambigator* sufficiently resolves the indexing ambiguity such that REFMAC no longer identifies the data as twinned.
